# Development of a PROTAC Targeting Chk1

**DOI:** 10.1101/2023.12.30.573733

**Authors:** Sandipan Roy Chowdhury, Patrick Chuong, Victoria E. Mgbemena, Alexander Statsyuk

## Abstract

A series of Chk1 degraders were designed and synthesized. The degraders were developed through the conjugation of a promiscuous kinase binder and thalidomide. One of the degraders PROTAC-2 was able to decrease Chk1 levels in a concentration-dependent manner in A375 cells. The developed probes can be useful for the development of selective and more potent Chk1 degraders.

---

Chk1 is activated by DNA damage (DNA damage checkpoint), and delays cell cycle progression through S and G2/M phases.^1–3^ Knockcout of both Chk1 alleles is lethal in mice, while heterozygous Chk1^+/-^ mice are healthy and are not predisposed to early tumorigenesis for up to 18 months.^4^ Further studies showed that Chk1 is required for embryonic stem (ES) cell proliferation, and Chk1 deficient ES had defects in proliferation accompanied by apoptotic cell death. Interestingly, Chk1^-/-^ embryos died from p53-independent apoptosis. Since p53 controls both the G2/M and the G1 cell cycle checkpoints,^5,6^ these results suggest that Chk1 inhibition may selectively potentiate the effect of DNA damaging agents in p53-deficient tumors. Indeed, this was observed in human-in-mouse tumor models of triple negative breast cancers. Combination of Chk1 inhibitor AZD7762 and topoisomerase I inhibitor irinotecan significantly extended the survival of tumour bearing mice and lead to substantial decrease in tumour volume, compared to those treated with either AZD7762 or Irinotecan, but not both.^7^ Recent studies have also shown that Chk1 inhibitors can potentiate the effect of immune checkpoint inhibitors on tumors that normally do not respond to immune therapy because they lack PD-L1 on their surface. In the case of small cell lung cancer, pharmacological inhibition of Chk1 led to the increased cytosolic dsDNA, leading to activation of STING pathway, and increased expression of PD-L1 on the cell surface. Treatment of H82 cell bearing mice with the combination of Chk1 inhibitor prexasertib and immune checkpoint inhibitor Atezolimumab have led to a complete abrogation of tumors in mice, to the point that it became difficult to find residual tumors for subsequent histology analysis.^8^

Given its promising properties as a drug target, numerous Chk1 inhibitors are undergoing preclinical and clinical trials.^9^ Traditional approaches to inhibit Chk1, center around ATP competitive inhibitors. However, this approach suffers from the lack of selectivity due to the highly conserved nature of the ATP binding site of protein kinases (~500 known). Moreover, it has been shown that Chk1 stabilizes and promotes replication fork progression, and this is not dependent on its enzymatic kinase activity.^10,11^ Targeted protein degradation is therefore an alternative approach for CHK1 inhibition. This can be achieved by hijacking the natural mechanism of Chk1 inactivation by degradation. Chk1 is normally activated by phosphorylation on Ser-345 by ATR kinase, but this phosphorylation also triggers Chk1 ubiquitination and degradation by numerous E3 ligases.^12,13^ A model was proposed that Chk1 degradation serves as a negative feedback loop to eventually turn Chk1 off to allow cell cycle progression. Indeed, it has been shown that topoisomerase I inhibitor camptothecin induces Chk1 phosphorylation and its subsequent degradation by <u>C</u>ullin-<u>R</u>ING E3 ligases (CRL).^13^ It was further shown that camptothecin resistant cell lines MDA-MB-231 and TK-10 have low expression of CRL component Fbx6, leading to a less efficient degradation of Chk1. Taken together it is reasonable to hypothesize that small molecule degraders of Chk1 will serve as effective substitutes of the natural mechanism of Chk1 inactivation. However, Chk1 degraders are not currently known.

PROTACs are bifunctional molecules that recruit E3 ligases to their protein targets to induce ubiquitination and degradation. Recently a series of PROTACs were prepared that contained promiscuous kinase binder attached to an E3 ligase recruiting moiety. Surprisingly, such PROTACS were selective degraders, due to the requirement for the ternary complex formation, and the accessibility of lysines for ubiquitination.^14,15^ Furthermore, such PROTACs can be used as a convenient scanning tool to quickly test if a particular kinase can be degraded by the particular E3 ligase in cells. Yet in all cases degradation of Chk1 was not observed, suggesting that Chk1 is not degradable at least by Cereblon or VHL E3 ligases.^15^ It is very well known however that single atom changes can make the difference between efficient and non-efficient protein degradation.^16^ We therefore decided to test additional protein degrader probes **PROTAC 1-4** in the hopes to find Chk1 degraders. Our model degraders consist of the previously reported promiscuous kinase inhibitor **1**, which is linked to the Cereblon recruiting moiety thalidomide *via* a triazole-PEG linker.^17^ We hypothesized that prepared compounds may serve as useful probes to scan degradable kinome for many kinases in the future. Aminopyrazole **1** (Fig. 1) contains pyrazolopyrimidine pharmacophore for promiscuous kinase inhibition.^18^ When screened against a panel of 359 wild-type kinases at 10 mM concentration, it was able to bind to 337 of them including Chk1.^17^ Therefore **1** can be used as a warhead to study the degradation of Chk1. Next, we wanted to identify an appropriate linkage position for the PEG linker such as to not disrupt the critical binding interactions between aminopyrazole **1** and the kinase. Crystal structure of the compound **2**, an analogue of compound **1**, with c-Src revealed that position 5 and 6 of **1** could be solvent exposed and can be used as a handle to attach polyethylene glycol (PEG) linker conjugated with thalidomide (Fig. 2).^17^ Based on the above analysis we decided to prepare a series of **PROTACs 1-4** by conjugating **1** to thalidomide via flexible PEG linkers of different lengths (Fig.3).

**Figure 1.**
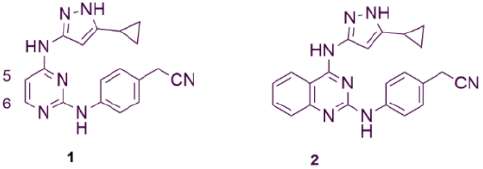
Chemical structure of promiscuous inhibitors.

**Figure 2.**
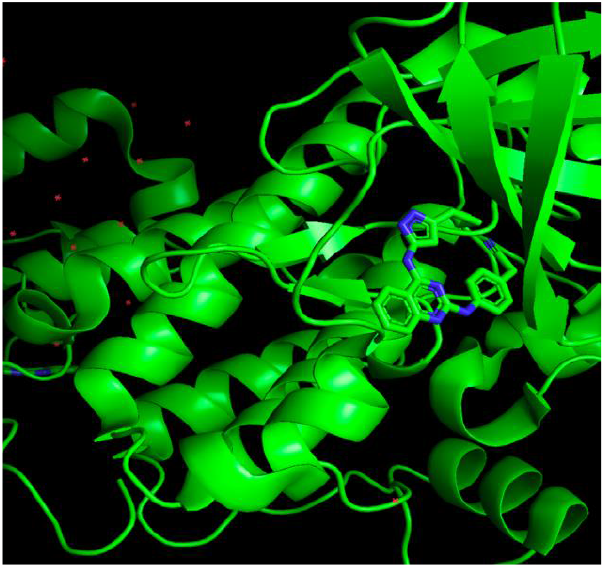
Crystal structure of **2** with c-src (PDB code-3F6X)

**Figure 3.**
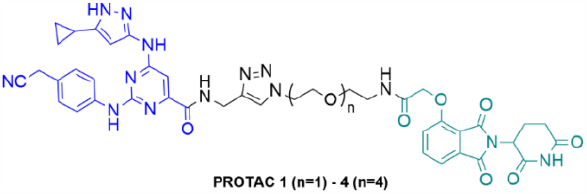
Chemical structure of the promiscuous kinase degraders.

The synthetic route to access PROTAC **1-4** is summarized in Scheme 1 and Schemes S1-S2. Briefly, commercially available Methyl 2,6-dichloropyrimidine-4-carboxylate and 3-Cyclopropyl-1H-pyrazol-5-amine were reacted via an S_N_Ar substitution reaction to form compound **3** (Scheme S1).^19^ A second nucleophilic substitution on **3** with 4-aminophenylacetonitrile yielded compound **5**. Subsequent hydrolysis of the methyl ester followed by coupling with propargyl amine led to compound **7**, which was coupled to thalidomide-PEG conjugates using CuAAC click chemistry.

We chose A375, a malignant melanoma cell line, as a model system to study PROTAC-mediated degradation of Chk1. First, we treated A375 cells with the different concentrations of the PROTAC **1-4** for 18 h and assessed degradation of Chk1 by Western blot analysis (Figure 4).

**Figure 4.**
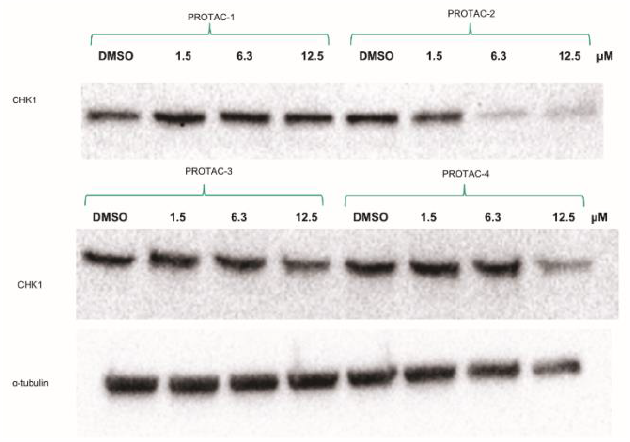
Effect of PROTAC 1-4 in A375 cells. Western blot analyses of Chk1 with lysates generated from A375 cells treated with different concentration of the PROTACs for 18 h at 37 °C and 5% CO2. α-Tubulin was used as a loading control.

Our findings suggested that **PROTAC-2** with a linker length of 18 atoms (n=2) was the most efficient Chk1 degrader whereas **PROTAC-1** was the least effective as a degrader among **PROTACs 1-4** (Figure 3). Although **PROTAC-3** and **4** were not as efficient as **PROTAC-2** but they caused some degree of Chk1 degradation at the highest concentration (12.5 μM).

**Scheme 1.**
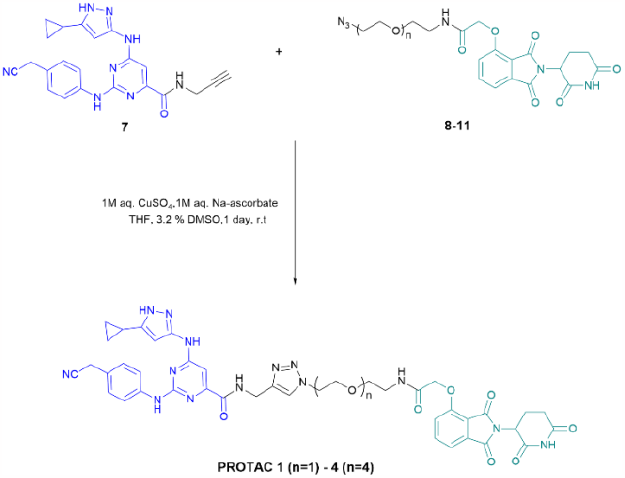
Synthesis of the PROTAC 1-4.

Inspired by these results, we next evaluated a dose-response study of **PROTAC-2**. A375 cells were treated with different concentrations of **PROTAC-2** for 12 h followed by quantification of Chk1 levels by Western blot (Fig. 5A). Our findings suggested that **PROTAC-2** degraded Chk1 with the DC_50_ (the concentration at which half-maximal degradation is observed) of 1.33 μM (Fig. 5B and 5C). Interestingly, we did not observe the hook effect for **PROTAC-2**, nor did we observe a complete degradation of Chk1, even at higher concentrations such as 12 μM. (Fig. 5). Chk1 degradation by PROTAC-2 was inhibited in the presence of MG132 and pomalidomide, suggesting that CHK1 degradation occurs via the proteasome and cereblon (Fig. 6).

**Figure 5.**
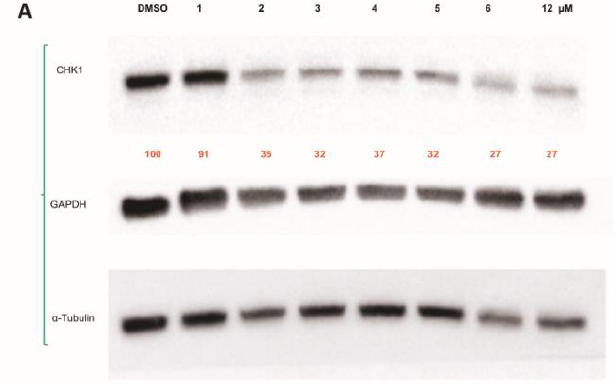
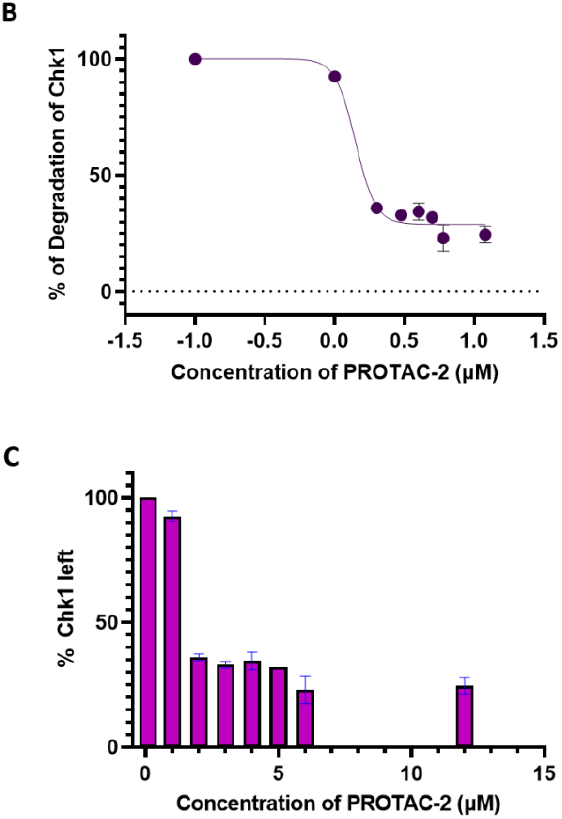
Dose-response curve for PROTAC-2. (A) Western blot analyses of CHK1 with lysates generated from A375 cells treated with different concentration of the PROTAC-2 for 12 h at 37 °C and 5% CO2. GAPDH and α-Tubulin was used as loading controls. (B) and (C) Percentage of CHK1 degradation and DC_50_ of PROTAC-2.

**Figure 6.**
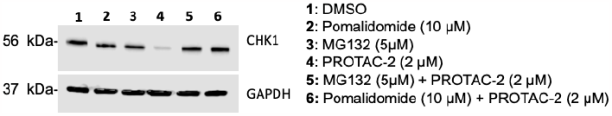
Treatment of A375 cells with PROTAC-2 or co-treatment of A375 cells with PROTAC-2 and MG132 or Pomalidomide. A375 cells were treated with the indicated concentration of PROTAC-2, MG132 or Pomalidomide for 20h and lysates were subjected to Western Blot analysis.

In summary, we reported a series of chemical probes to explore degradable kinome and applied these probes to test the degradation of Chk1 kinase by E3 ligase Cereblon. PROTAC-2 was a potent Chk1 degrader with DC_50_ of 1.33 µM. In principle, similar chemistry can be used to conjugate aminopyrazole **1** with other E3 ligase recruiting moieties to test their ability to degrade Chk1. Further development of selective Chk1 degraders and their use as anticancer therapeutics will be reported in the future.

## Supporting information

Experimental methods

## Acknowledgment

Research reported in this publication was supported by the National Institute of General Medical Sciences of the National Institute of Health under Award Number R01GM115632. The content is solely the responsibility of the authors and does not necessarily represent the official views of the National Institute of Health. The authors would like to thank Uyino Victory Eromosele for helping with producing the WB for Figure 6.

