## Supplementary material for "Development of a PROTAC Targeting Chk1": Experimental methods

1. Chemical synthesis
a. General methods……………………………………………………………………………………………………………3
b. Synthetic procedures……………………………………………………………………………………………………
Synthesis of Chk1 ligand - Scheme S1…………………………………………………………………………
Synthesis of thalidomide-linker conjugate Scheme S2…………………………………………………
Synthesis of PROTAC **1-4**……………………………………………………………………………………………
Mass spectra…………………………………………………………………………………………………………………
NMR spectra…………………………………………………………………………………………………………………
2. Biological methods……………………………………………………………………………………………………………9
a. General methods………………………………………………………………………………………………………………
b. Experimental procedures…………………………………………………………………………………………………
Testing of PROTAC **1-4**……………………………………………………………………………………………………
Proteasome dependency and competition experiment……………………………………………………

3. Reference………………………………………………………………………………………………………………………10

### 1. Chemical Synthesis

**a. General Experimental Methods.** All experiments requiring anhydrous conditions were conducted in flame-dried glassware fitted with rubber septa under a positive pressure of dry nitrogen. Column chromatography was performed on a Combiflash $R_f$ + system using prepacked silica gel columns from Silicycle. An acetone cooling bath was adjusted to the appropriate temperature by the addition of small portions of dry ice.

**Instrumentation.** HRMS were obtained in a ThermoScientific LUMOS Tribid mass spectrometer. HPLC purification was conducted with an Agilent 218 Injection pump coupled with a Varian-ProStar 325 detector and a Restek Pinnacle DB $C_{18}$ column (250 × 21.2 mm, 5 µm). $^1$H NMR was obtained on a Varian (Paolo Alto, CA) 400 MR spectrometer. The following abbreviations were used to explain the multiplicities; d = doublet, s = singlet, br = broad.

**Materials.** Reagents were purchased from Aldrich Chemical Co., Sigma Chemical Co. or Combi Blocks and were used without further purification. Azido-PEG-amines were bought from BroadPharm. **X** was prepared following previous literature procedures.[1]

### b. Synthetic procedures

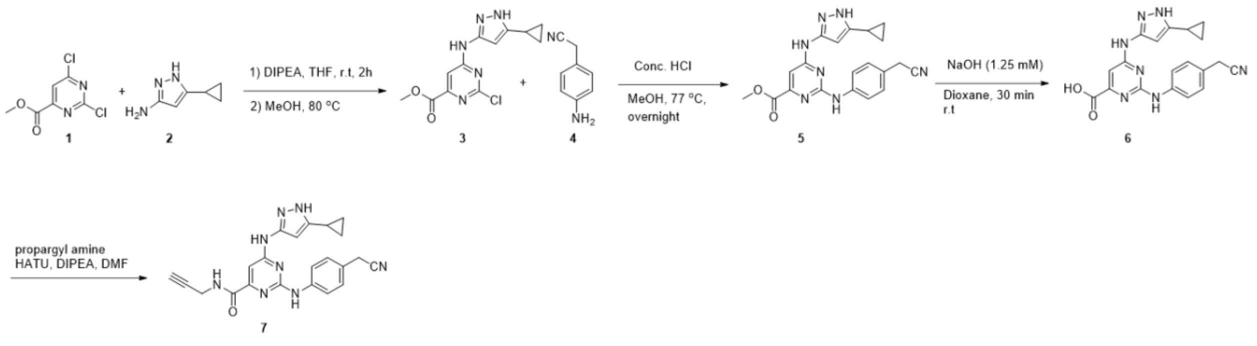


**Scheme S1**. Synthesis of the Chk1 ligand **7**

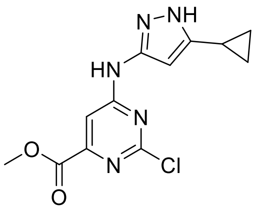


**3**

**Methyl 2-chloro-6-((5-cyclopropyl-1*H*-pyrazol-3-yl)amino)pyrimidine-4-carboxylate (3)**. To a solution of Methyl 2, 4-dichloropyrimidine-4-carboxylate (**1**, 12.1 mmol) in 15 mL anhydrous THF at 0 °C was added 3-Amino-5-cyclopropyl-1*H*-pyrazole (**2**, 12.2 mmol) and DIPEA (3.74 mmol) in 15 mL anhydrous THF and the reaction mixture was stirred at r.t for 2 h. The solvents were evaporated in vacuo, and the residue was heated in methanol (15 mL) at 65 °C for 1 h. The mixture was cooled to r.t and filtered. The precipitate was dried overnight to obtain a yellow powder: yield 2.53 g (71%)

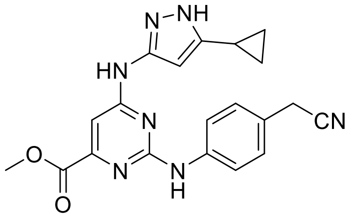


**5**

**Methyl 2-((4-(cyanomethyl)phenyl)amino)-6-((5-cyclopropyl-1H-pyrazol-yl)amino)pyrimidine-4-carboxylate (5)**. To a suspension of **3** (0.5 g, 1.7 mmol) and 2-(4-aminophenyl)acetonitrile (**4**, 0.22 g, 1.7 mmol) in 6 mL MeOH was added 0.3 mL of conc. HCl. The reaction mixture was heated at 93 °C overnight. The reaction mixture was cooled to r.t and filtered. The yellow precipitate was dried to obtain **5** as a yellow solid: yield 0.34 g (51%).

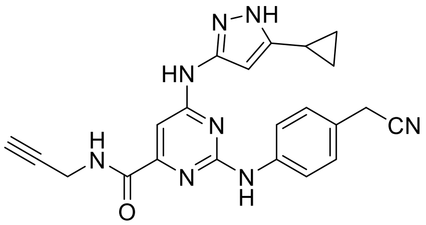


**7**

**2-((4-(cyanomethyl)phenyl)amino)-6-((5-cyclopropyl-1H-pyrazol-3-yl)amino)-N-(prop-2-yn-1-yl)pyrimidine-4-carboxamide** (**7**). To a stirred solution containing 50 mg of **5** in 5 mL of 2:1:2 MeOH, 1,4-dioxane and water at 77 $^oC$ was added 1 M NaOH dropwise and stirred until TLC indicated complete consumption of starting material. The intense yellow mixture was cooled to r.t and acidified with 2 M HCl until pH ~ 2, a white precipitate appeared. The mixture was concentrated. The crude was again suspended in 1:1 water-acetonitrile and lyophilized to obtain a yellowish white powder. The crude **6** was used in the next step without further purification.

To the crude **6** was added 2 mL of anhydrous DMF followed by DIPEA (80 μl) and HATU (64 mg) at r.t. The suspension was stirred at r.t for 15 min and propargyl amine (10 mg) was added dropwise. The mixture was stirred at r.t overnight. The reaction mixture was diluted with 50 mL ice-cold water. The aqueous layer was extracted with two 25-mL portions of ethyl acetate. The combined organic layer was dried over anhydrous $MgSO_4$, filtered and concentrated. The crude was suspended in 10 mL of ethyl acetate and filtered to obtain **7** as a yellowish white solid: yield 28 mg (52%). $^1H$ NMR (600 MHz, DMSO-$d_6$) $\delta$ 0.64 (br. s, 2H), 0.89 (br. s, 2H), 1.19 (s, 1H), 1.84 (br. s, 1H), 3.94 (s, 2H), 4.06 (s, 2H), 7.23 (d, $J$ = 12 Hz, 2H), 7.69 (br. s, 2H), 8.39 (br. s, 1H), 9.29 (br. s, 1H), 12.08 (br. s, 1H). **MS** (ESI) m/z: calculated for $C_{22}H_{21}N_8O^+$, 413.2; found 413.3.

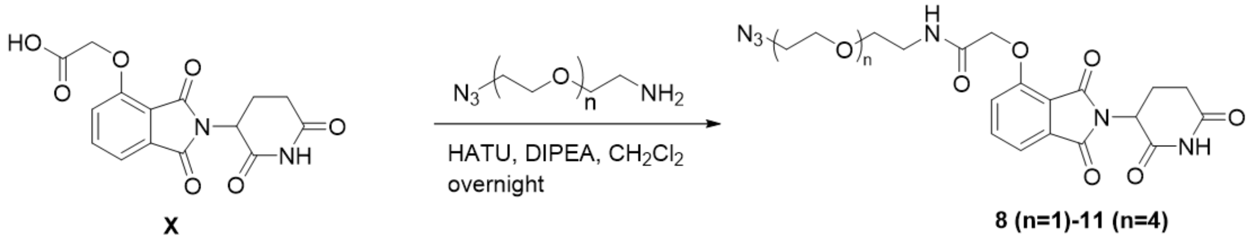


**Scheme S2**. Synthesis of the thalidomide-linker conjugate **8**-**11**

To a stirred suspension containing 2-((2-(2,6-dioxopiperidin-3-yl)-1,3-dioxoisoindolin-4-yl) oxy) acetic acid (X) in anhydrous $CH_2Cl_2$ was added DIPEA (3 eq.) and HATU (1.1 eq.) respectively. The reaction mixture was stirred at r.t for 15 min and Azido-PEG$_n$-amine (1.1 eq.) was added dropwise. The yellow reaction mixture was stirred overnight at r.t under nitrogen atmosphere. The reaction mixture was diluted with 10 mL of ice-cold water. The aqueous layer was extracted with three 10-mL portions of ethyl acetate. The combined organic layer was dried over anhydrous $MgSO_4$, filtered and concentrated under diminished pressure. The residue was purified first on a silica gel column using dichloromethane-methanol followed by on $C_{18}$ reverse phase HPLC using a gradient of 0% to 100% acetonitrile in water over a period of 20 min and lyophilized.

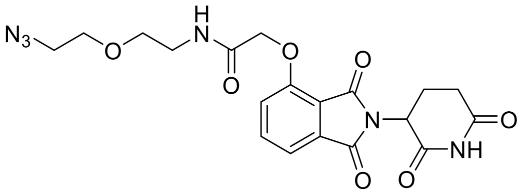

**8**

Azido-PEG$_1$-thalidomide (**8**): yield 30 mg (50%)

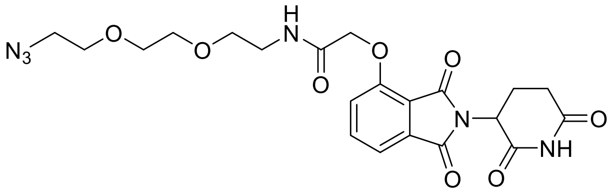

**9**

Azido-PEG$_2$-thalidomide (**9**): yield 20 mg (46%)

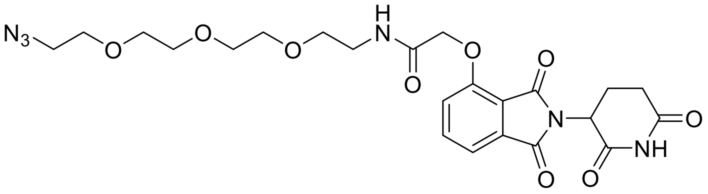

**10**

Azido-PEG$_3$-thalidomide (**10**): yield 27 mg (60%)

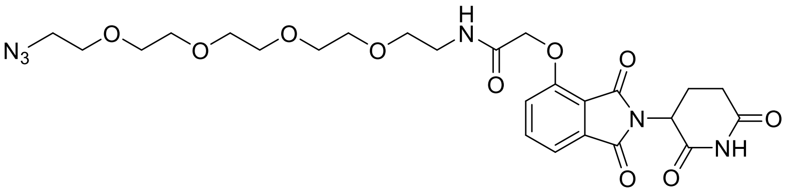

**11**

Azido-PEG$_4$-thalidomide (**11**): yield 26 mg (67%)

**Synthesis of PROTAC 1-4**

**7** (1 eq.) and **8-11** (1.2 eq.) were dissolved in 0.5 mL THF containing 16 ul of DMSO. To this solution were added 15 µL of 1 M $CuSO_4$ followed by 15 µL of 1 M Na-ascorbate. The reaction mixture was stirred for 1 h at r.t. An additional 15 µL of both $CuSO_4$ and Na-ascorbate were added to the reaction mixture and stirred for 1 h. The completion of the reaction was monitored by $C_{18}$ TLC using 1:1 water-acetonitrile as the mobile phase. The reaction mixture was concentrated under diminished pressure. The crude was re-dissolved in 1:1 water-acetonitrile and purified on $C_{18}$-reversed phase HPLC (250 x 21.2 mm) using a gradient of 0 to 100% acetonitrile in water over a period of 20 min. The desired fractions from 16.5 to 17.5 min were collected and lyophilized to obtain white solids.

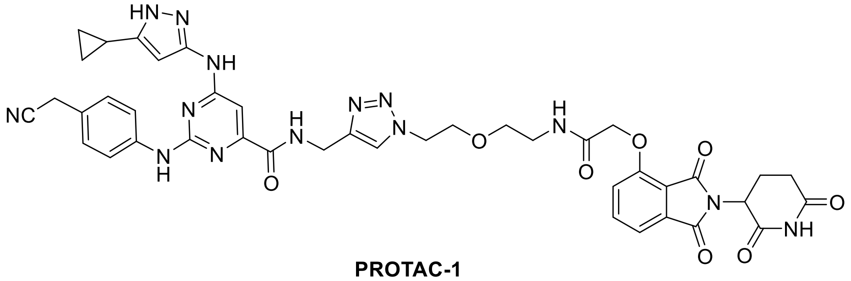

**PROTAC-1**

Yield-6.0 mg (40%); **HRMS** (ESI) m/z: calculated for $C_{41}H_{42}N_{14}O_8^{2+}$, 429.1650; found 429.1648.

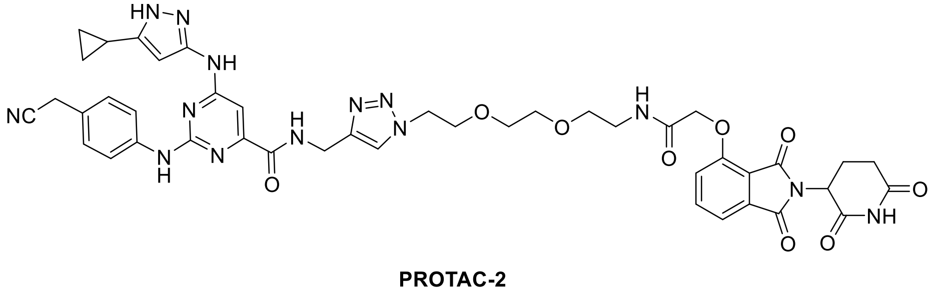

**PROTAC-2**

Yield-5.7 mg (35%); **HRMS** (ESI) m/z: calculated for $C_{43}H_{46}N_{14}O_9^{2+}$, 451.1781; found 451.1772.

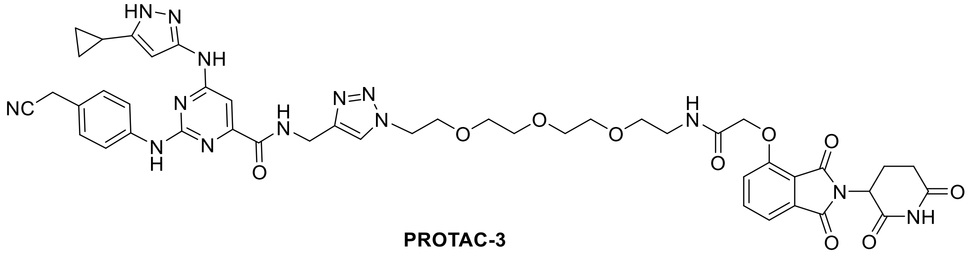

**PROTAC-3**

Yield-7.0 mg (37%); **HRMS** (ESI) m/z: calculated for $C_{45}H_{50}N_{14}O_{10}^{2+}$, 473.1912; found 473.1906.

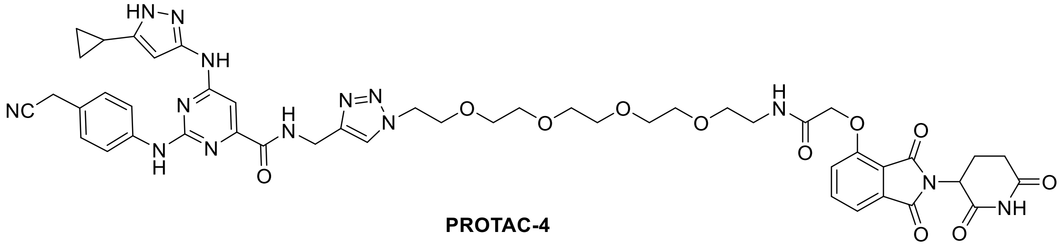


**PROTAC-4**

Yield-5.5 mg (28%); **HRMS** (ESI) m/z: calculated for $C_{47}H_{54}N_{14}O_{11}^{2+}$, 495.2043; found 495.2034.

### Mass spectra for PROTACs

### PROTAC-1

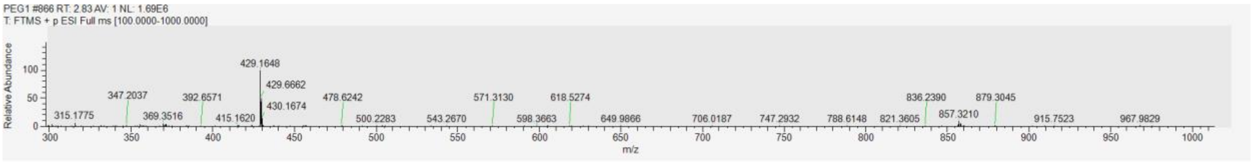


### PROTAC-2

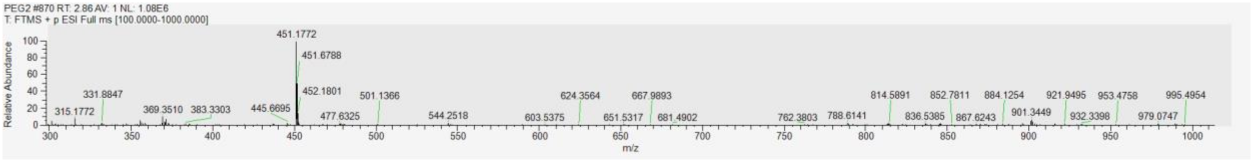


### PROTAC-3

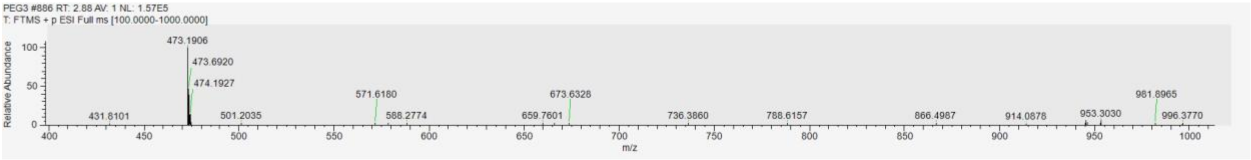


### PROTAC-4

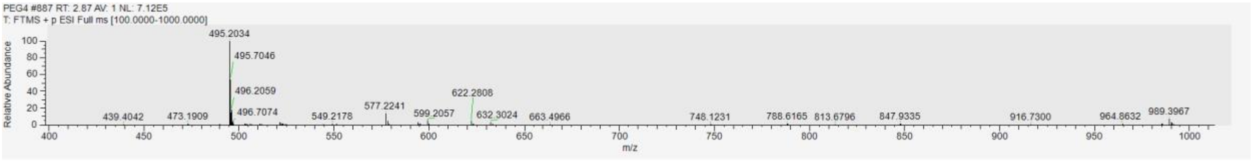

**[1]H NMR Spectra**

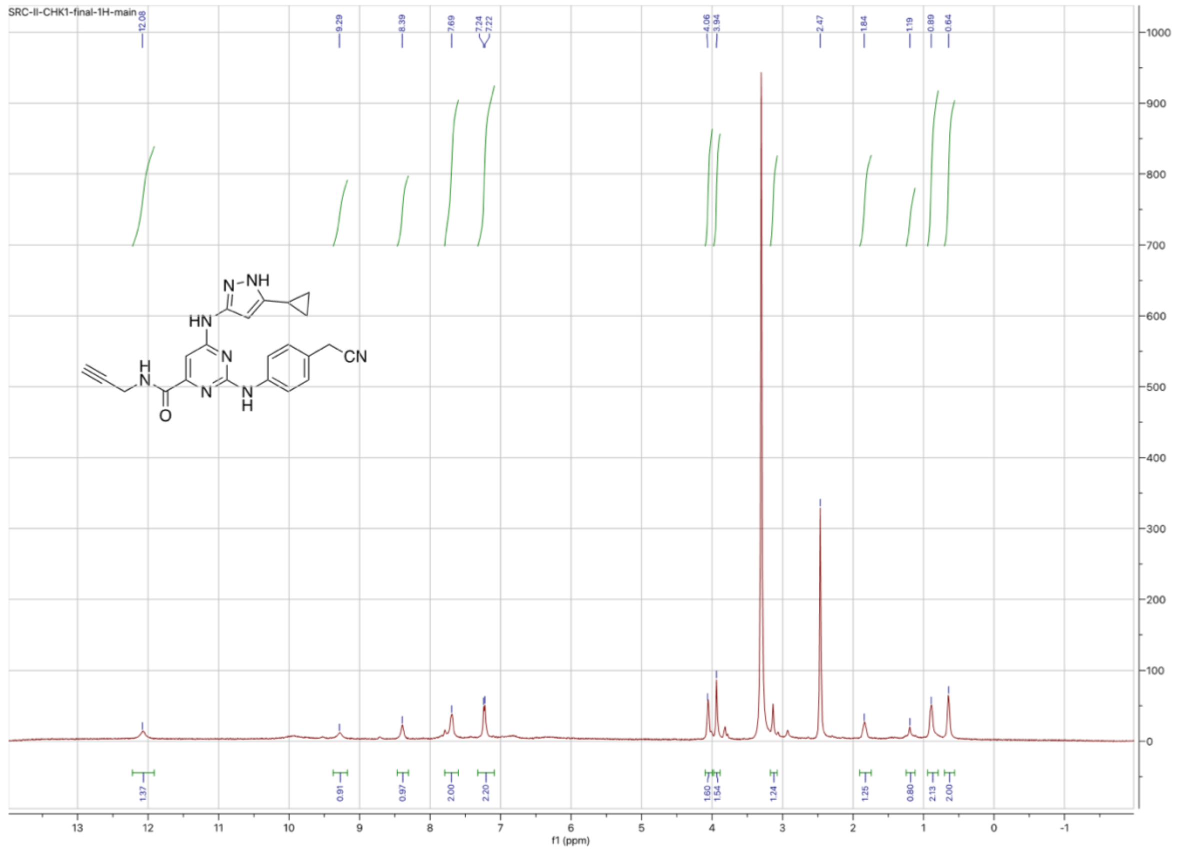

### 2. Biological methods

### a. General methods

#### Reagents

All reagents were purchased from commercial sources and used as supplied unless otherwise indicated. RIPA lysis buffer were purchased from Thermo Fisher Scientific. A375 cells, DMEM media and heat deactivated fetal bovine serum (FBS) were purchased from ATCC. Complete Mini EDTA free Protease inhibitor cocktail was purchased from Sigma.

**Cell culture**

A375 cells were cultured in Dulbecco's Modified Medium (DMEM, ATCC) supplemented with 10% heat-inactivated fetal bovine serum (FBS, ATCC). All cells were maintained in a humidified incubator at 37 $^oC$ and 5% $CO_2$. Cells were seeded on plates at least 24 h before experiment.

**Western blot analysis of drug treated cells.**

A375 cells (melanoma cell line) were plated at 6 x $10^5$ cells in 60-mm dishes and treated with: 5µM MG132, PROTAC-**1-4**, 10 µM Pomalidomide, or DMSO. Collected samples were lysed in 1X RIPA Buffer with protease inhibitor cocktail (18-24h post treatment), sonicated, centrifuged, and stored at -80 degrees Celsius if needed. 30$\mu$g of cell lysates per well were loaded to SDS-PAGE gel, followed by western blotting.

Cell signaling antibodies Primary: 1:1000. Secondary 1:5000.
CHK1: Cell signaling #37010S, E9M4D, Rabbit mAb.
GAPDH: Cell signaling #5174S, D16H11 Rabbit mAb.
$\alpha$-tubulin: Cell signaling 11H10, Rabbit mAb #2125.
Signal detection: Signal Fire ECL reagent (CST # 6883).
Signal imaging: BioRad Chemidoc system.
